# Ancient hybridization and strong adaptation to viruses across African vervet monkey populations

**DOI:** 10.1101/088989

**Authors:** Hannes Svardal, Anna J Jasinska, Cristian Apetrei, Giovanni Coppola, Yu Huang, Christopher A Schmitt, Beatrice Jacquelin, Michaela Müller-Trutwin, George Weinstock, J Paul Grobler, Richard K Wilson, Trudy R Turner, Wesley C Warren, Nelson B Freimer, Magnus Nordborg

## Abstract

Vervet monkeys (genus *Chlorocebus*, also known as African green monkeys), are highly abundant in savannahs and riverine forests throughout sub-Saharan Africa. They are amongst the most widely distributed nonhuman primates, show considerable phenotypic diversity, and have long been an important biomedical model for a variety of human diseases^1^ and in vaccine research^2–4^. They are particularly interesting for HIV/AIDS research as they are the most abundant natural hosts of simian immunodeficiency virus (SIV), a close relative of HIV. Here we present the first genome-wide survey of polymorphism in vervets, using sequencing data from 163 individuals sampled from across Africa and the Caribbean islands where vervets were introduced during the colonial era. We find high diversity, within and between taxa, and clear evidence that taxonomic divergence was reticulate rather than following a simple branching pattern. A scan for diversifying selection across vervet taxa yields gene enrichments much stronger than in similar studies on humans^5^. In particular, we report strong and highly polygenic selection signals affecting viral processes — in line with recent evidence that proposes a driving role for viruses in protein evolution in mammals^6^. Furthermore, selection scores are highly elevated in genes whose human orthologs interact with HIV, and in genes that show a response to experimental SIV infection in vervet monkeys but not in rhesus macaques, suggesting that part of the signal reflects taxon-specific adaptation to SIV. Intriguingly, rather than affecting genes with antiviral and inflammatory-related functions^7^, selection in vervets appears to have primarily targeted genes involved in the transcriptional regulation of viruses, and in particular those that are harmful only under immunodeficiency, suggesting adaptation to living with SIV rather than defense against infection.

## Main

The genus *Chlorocebus* has been viewed as a single species (*Ch. aethiops*) with several subspecies or as 5-6 species with additional subspecies^8^. Our sample was intended to capture this diversity (Supplementary Data 1), by including individuals from nine countries in Africa and two countries in the West Indies, into which vervets were introduced in the 1600s, and subsequently established sizable feral populations (Fig. 1A, Supplementary Data 1). No previous study has conducted genome-wide resequencing in a non-human primate in such a large sample and over such a geographically extensive area. Employing a standard pipeline for alignment to the reference *Chlorocebus_sabaeus* 1.1^9^, derived from a St. Kitts-origin monkey, and joint variant detection across all samples, we discovered a total of over 97 million single nucleotide polymorphisms (SNPs), 61 million of which passed our quality filters (Online Methods, Supplementary Figs. 1 to 3).

**Fig. 1:**
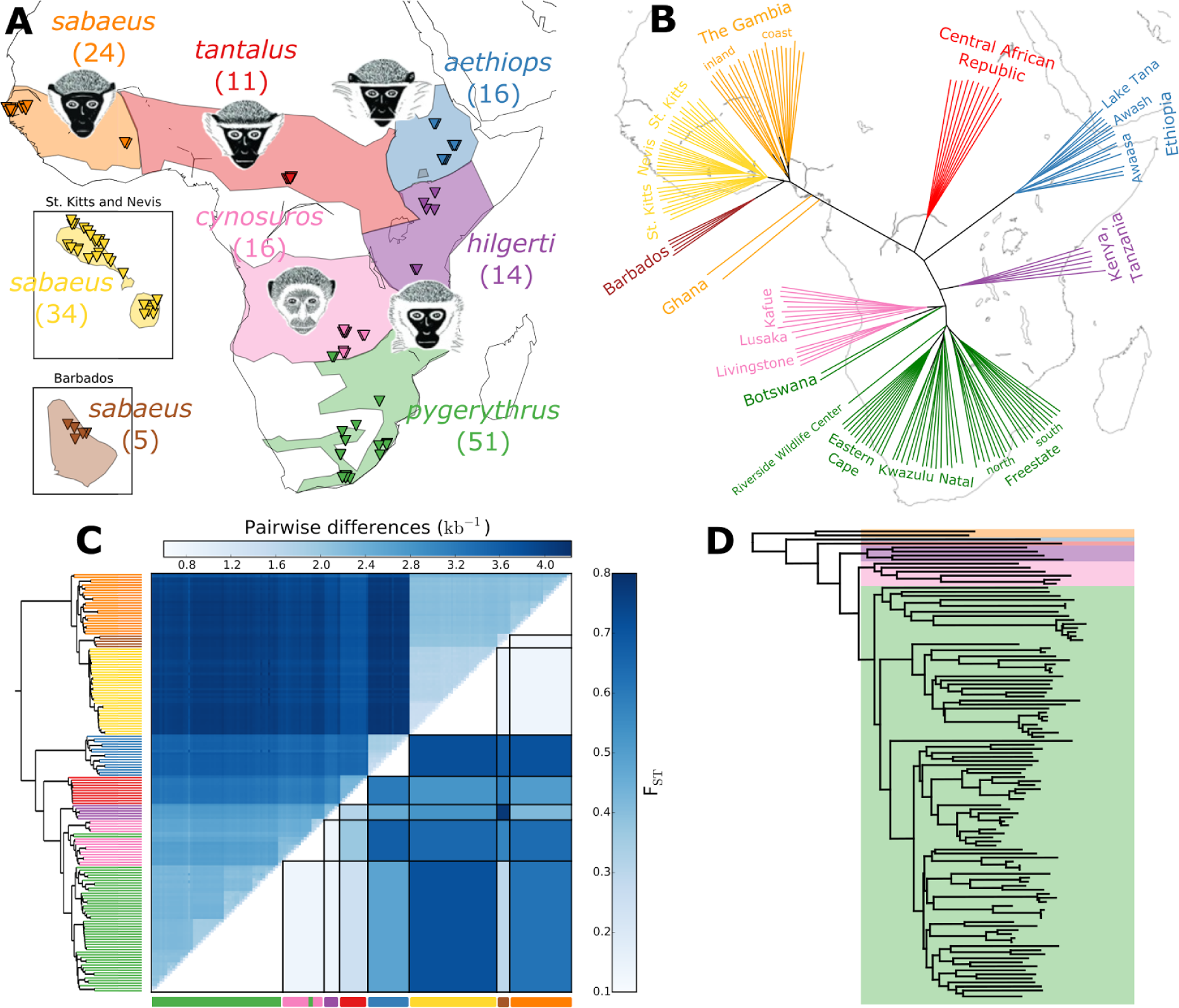
Sample information and genetic relatedness. (A) Taxon-distribution, approximate sampling locations (triangles) and representative drawings (where available: *hilgerti* and *pygerythrus* are morphologically very similar and have often not been considered separately). Number of whole-genome sequenced monkeys given in parenthesis. (B) Neighbor-joining tree based on pairwise differences oriented to approximately fit geographic sampling locations. (C) Matrix-plot of pairwise genetic differences per callable site (above diagonal) and fixation index (F_ST_) between groups (below diagonal). Rows and columns are sorted according to a hierarchical clustering (UPGMA) tree of vervet pairwise genetic differences. (D) Clustering tree of SIVagm pol gene sequences sampled from wild vervets (Africa only). Tree was constructed with sequences from both vervets included in this study and with sequences from the HIV Sequence Databases. The branches have been colored to match the group labels in (A).

Clustering of individuals based on pairwise genetic distance (Figs. 1B and 1C) generally agrees with prior morphological and geographic classification, and led us to define six African and two Caribbean taxonomic groups: *sabaeus* (West Africa), *aethiops*, *tantalus*, *hilgerti*, *cynosuros*, *pygerythrus*, *sabaeus* (St.Kitts and Nevis) and *sabaeus* (Barbados). Genetic relatedness also suggests isolation-by-distance within groups (Fig. 1B). Both geographic location and group identity contribute significantly to explaining the overall pattern of polymorphism (likelihood-ratio test p<2^*^10^−3^ and p<10^−52^, respectively, Supplementary Table 3). Our data agree with the morphology-based taxonomy^8^ in that *sabaeus*, *aethiops* and *tantalus* appear to be well-defined taxa, whereas *hilgerti*, *cynosuros* and *pygerythrus* are comparatively closer to each other, exhibiting substantial amounts of shared variation and strongly correlated allele frequencies (Fig. 1C, Supplementary Fig. 4). However, while morphological evidence groups *hilgerti* and *pygerythrus* as a single species (*Ch. pygerythrus*) distinct from *cynosuros* (*Ch. cyonosuros*), our data show that *pygerythrus* and *cynosuros* are closer to each other than either is to *hilgerti*. Indeed, two *pygerythrus* individuals from Botswana are more closely related to *cynosuros* than to other *pygerythrus*. This is probably due to admixture: as we note below, there is abundant evidence for admixture between these groups. Finally, the pattern of relatedness among SIV strains mirrors the pattern in vervets (Fig. 1D), suggesting that SIV existed in vervets prior to their initial divergence more than half a million years ago and has co-evolved with the taxa^9,10^.

Our data also clarify the origin of the Caribbean vervets. While it has long been known that these populations are derived from Western African *sabaeus*, it has not been clear how they are related to each other. The fact that vervets from Barbados are as different from vervets from St. Kitts and Nevis as they are from Gambian *sabaeus* (Fig. 1C) suggests that these two Caribbean populations were founded independently and experienced two independent bottlenecks (28% and 17% reduction in diversity relative to Gambia, respectively). The vervet population from Nevis, on the other hand, is genetically a subset of the St. Kitts population (17% reduction in diversity relative to St. Kitts), and likely was founded by individuals from this island, which is less than 4 km away. In human genetics there is currently great interest in sequencing studies of bottlenecked populations, both for elucidating population genetic processes and for identifying deleterious variants with a strong impact on phenotypes that have reached high frequency through drift. Caribbean vervets, founded through three distinct bottlenecks, may have unique value for such studies.

Returning to Africa, variation within vervet taxa is much larger than in humans^11^ and other great apes^12^ but is typical for other primates (Fig. 1C, Supplementary Fig. 5)^13^ — which is perhaps surprising given the ubiquity of vervets. Divergence between vervet taxa is generally higher than between subspecies of other primates, with average pairwise sequence divergence between taxa of ~0.4%, compared to 0.2% to 0.32% across great ape subspecies^12^, and F_ST_-values from 25% to 71% (Fig. 1C, Supplementary Fig. 6), compared to <15% across human populations^14^ or macaque subspecies^15^. However, maximum sequence divergence is substantially lower than between human and chimp (~1.24%)^16^. This intermediate status is supported by the presence of substantial amounts of both shared variation and fixed differences between vervet taxa (Supplementary Figure 4).

The process that gave rise to the current taxa was much more complex than a series of population splits. We used Admixture^17^ to cluster individuals into groups. This analysis generally resolves the above-mentioned taxa, and confirms the complicated relationships of south‐ and east-African vervets (Fig. 2A, Supplementary Figs. 7 and 8). However, there is evidence of admixture throughout. For example, Ghanaian *sabaeus* and Kenyan and Tanzanian *hilgerti* show substantial proportions of *tantalus* ancestry (Fig. 2A). D-statistics (ABBA-BABA test; Fig. 2, D and E)^18,19^ confirm that this shared ancestry represents gene flow or ancestral population structure between *tantalus* and both Ghanaian *sabaeus* and *hilgerti* (D = 15.2% and 5%, respectively; block jack-knifing p<10^−300^). Using the multiple sequentially Markovian coalescent method (MSMC; Figs. 2B and 2C, Supplementary Figs. 9 and 10., Supplementary Note 1)^20^ we show that the observed signatures of gene flow likely reflect ancient rather than recent admixture; for example, for *tantalus*, after initial divergence, gene flow ceased earlier with geographically more distant Gambian *sabaeus* than with geographically closer Ghanaian *sabaeus* (Fig. 2B). The lack of long shared haplotypes across taxa also supports the absence of recent admixture (Supplementary Table 3).

**Fig. 2:**
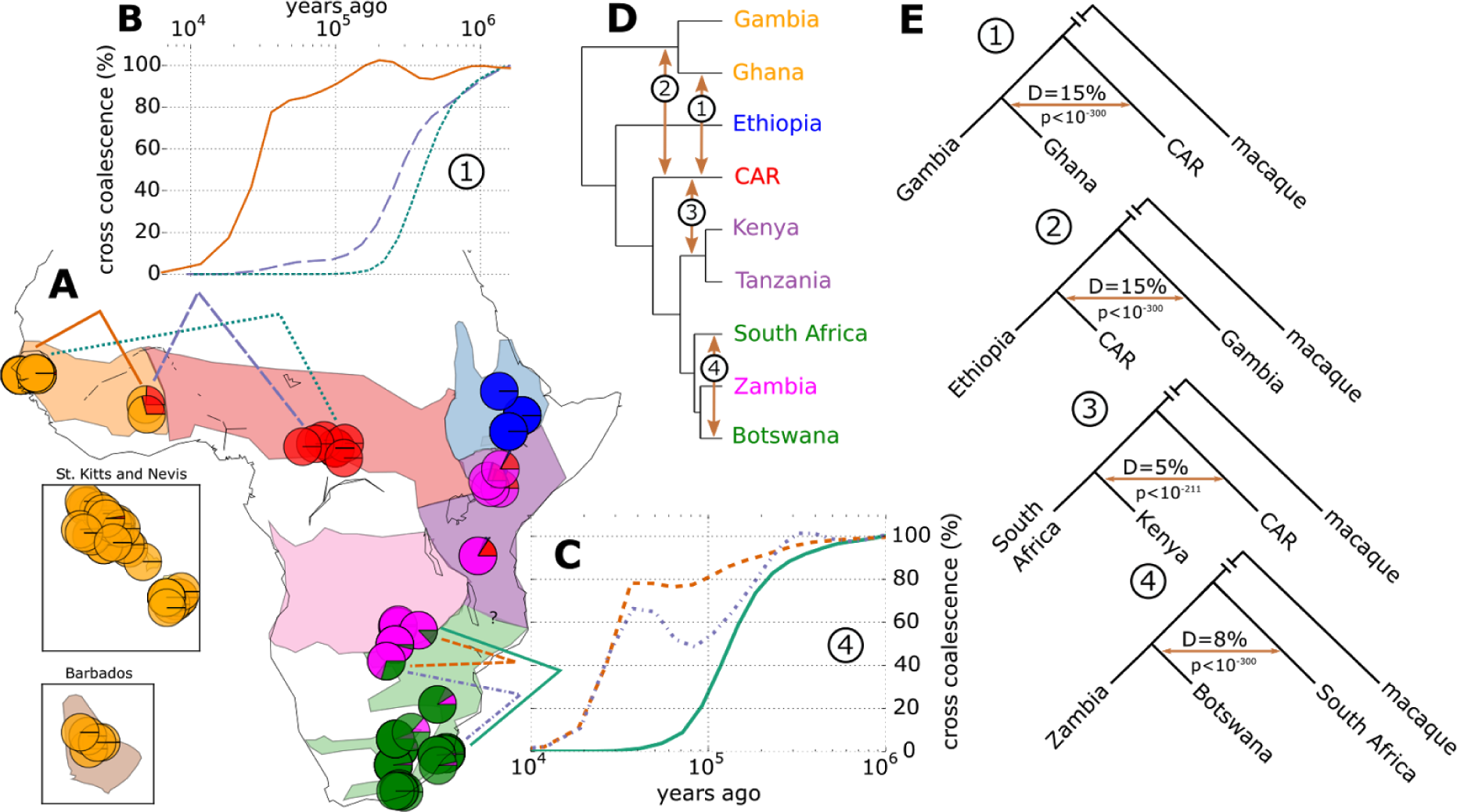
Evidence for gene flow across taxa. (A) Admixture clustering of individuals. Each pie-chart represents an individual and colours represent contributions from five assumed admixture clusters. Colored lines mark comparisons in panels B and C. (B) and (C) MSMC plots of cross-coalescence rate, a measure of gene flow, across time (on a log scale). (D) UPGMA tree of pairwise distance matrix summarized by country. Arrows point to evidence of cross-taxon gene flow. (E) D-statistics (ABBA-BABA test) for instances of gene flow shown in D. For full results see Supplementary Fig. 11 and Supplementary Data 2.

Turning to the east and south African *cynosuros*/*hilgerti*/*pygerythrus* complex, we find that, while simple clustering suggests that *hilgerti* is an outgroup to *cynosuros* and *pygerythrus* (Figs. 1B and 1C), Admixture represents *cynosuros* individuals as a mixture of *hilgerti* and *pygerythrus* with a larger contribution of the former (Fig. 2A). MSMC suggests a complex history of varying gene flow between the three groups (Supplementary Fig. 10). We also investigated the status of *pygerythrus* from Botswana, which appear as sister-group to Zambian *cynosuros* in the clustering tree (Fig. 2D): D-statistics shows that they have an additional genetic contribution from South African *pygerythrus* (Fig. 2E, D = 7.6%, jack-knifing p<10^−300^) and MSMC confirms an intermediate status of Botswanian *pygerythrus* with comparable levels of genetic exchange with both South African *pygerythrus* and *cynosuros* until total separation from both groups ten thousand years ago (Fig. 2C), again compatible with isolation by distance. In summary, our results are generally consistent with current taxonomy, but do not support the notion that these taxa evolved by a process of population splitting that would give rise to a clean phylogenetic tree. Instead, we see strong signatures of ancient gene flow and the overall pattern is one of gradual divergence and isolation by distance.

Our data provide a rare opportunity to look for signals of adaptation on a continent-wide scale, across multiple taxa. To identify footprints of selection, we used an approach that incorporates information on both the distortion of allele frequency spectra within groups and the increase in differentiation among pairs of groups at loci close to a group-specific selective sweep (XP-CLR)^21^. To summarize the 30 XP-CLR-comparisons between African taxa (Supplementary Figs. 12 to 19), we calculated “selection scores” — the root mean square XP-CLR scores (across taxon comparisons) — on a 1000 base pair grid along the genome.

These scores clearly capture strong signals of selection, because they are significantly higher in genic than intergenic regions (one-sided Mann-Whitney U test p<10^−300^, Supplementary Fig. 20). To gain further insight, we compared the distribution of average selection scores for genes across gene ontology (GO) terms (Supplementary Data 3) with the R-package TopGO^22^. Testing for enrichment using the relative rank of all scores yielded stronger signals than testing genes with the highest scores against the background, suggesting that weaker, polygenic effects contribute strongly to the signal of selection (Supplementary Fig. 21). We found 167 significantly enriched GO-terms, many of which are related to RNA transcription and cell signaling (Fig. 3A, Supplementary Fig. 22, Supplementary Data 4). These GO-enrichments show partial overlap with similar scores comparing human populations (Supplementary Fig. 23)^5^. However, enrichment p-values for vervet scores are many orders of magnitude more significant than their human counterparts suggesting that vervet taxa provide a powerful model to study diversifying selection across closely related primate taxa. The strongest selection scores are consistent with a dominant role of viral pathogens as selective agents in vervets. In particular, we note viral process (p<10^−9^), and positive and negative regulation of transcription from the polymerase II promoter (p<5^*^10^−17^, 5^*^10^−14^), which is known to interact with viral proteins (for example the HIV Tat gene during transcription elongation of HIV-1 LTP^23^). We note that evolutionary conservation scores^24^ do not show significant enrichment for the same virus-related gene categories (Supplementary Fig. 23), providing evidence that these signals are not predominantly driven by purifying selection (background selection), which can lead to confounding signals^25^.

**Fig. 3:**
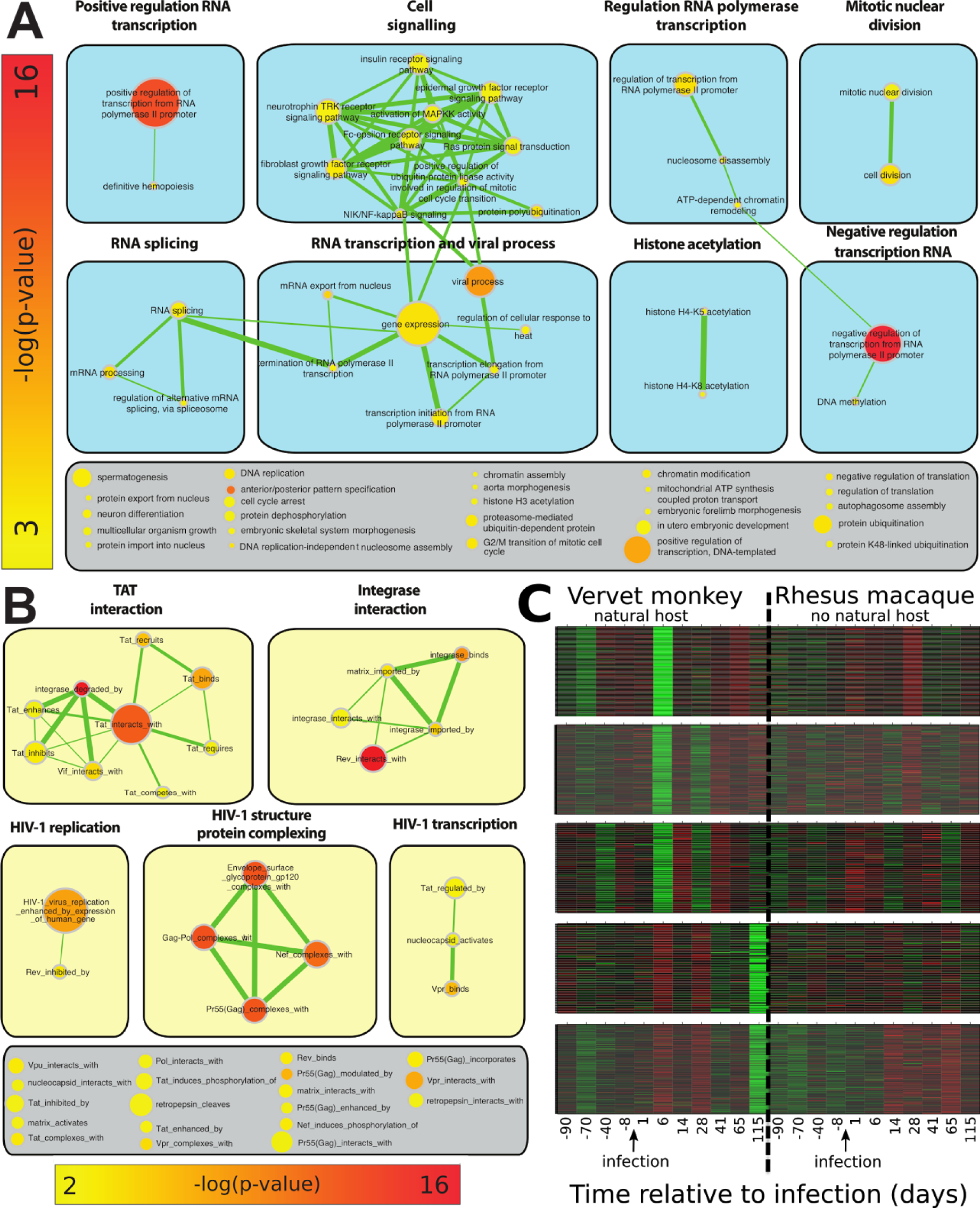
Gene set enrichment of signals of diversifying selection in the vervet monkey genome. (A) and (B) Enrichment-map network of categories which high average gene selection scores. Edges represent overlap in genes. Colors represent p-values on a log scale (red most highly significant). Node size represents number of genes in this category. Terms are grouped using Cytoscape clustermaker26. (A) Enrichment-map network of TopGo enriched GO terms with p < 0.001 (min genes=7; max=870). (B) Enrichment-map network of HIV-human interaction gene sets enriched in high mean selection scores with p<0.01 and Bonferroni-Holm FWER < 0.05 using a sumstat enrichment test (min genes=10; max=474). (C) Co-expression modules of genes differentially expressed in vervet or macaque pre‐ and post-SIV-infection that are also significantly enriched for high selection scores (FWER<0.05). Differentially expressed genes were grouped into co-expression modules using WGCNA. Of 33 gene expression modules from CD4+ blood cells, these five modules with significant enrichment for high selection scores are strongly biased towards vervet specific expression changes (Supplementary Fig. 28). Enrichment p-values for the represented modules are 10^−5^, 3.9^^*^^10^−4^,1.2^^*^^10^−3^, 10^−3^, 6.5^^*^^10^−3^ (top to bottom).

To test more specifically for virus-related selection signals, we looked for enrichment of signals among the orthologs of human HIV-interacting genes. Of the 166 sets of the NCBI HIV-1 human interaction database^27^ with more than ten genes in the vervet annotation, 45 show significant enrichment (Holm-Bonferroni familywise error rate (FWER) < 0.05, Fig. 3B, Supplementary Data 5). Not only are selection scores massively elevated in HIV-1 interaction gene sets compared to random background gene sets, they also are more highly elevated in HIV-1 categories than in regular GO-categories (Supplementary Fig. 25; one-sided Kolmogorov-Smirnov test, p<6^*^10^−5^), suggesting a dominant role for viruses in selection.

SIVagm is prevalent in African vervets^10,28^ and has diverged into taxon-specific strains (Fig. 1D) ^29,30^. Furthermore, while SIVagm is highly pathogenic when used experimentally to infect pigtailed macaques that are not natural SIV hosts^31,32^, infected vervets generally do not progress to AIDS, suggesting coevolution of virus and host. We hypothesized that coevolution between taxon-specific SIV strains and vervet taxa could lead to an ongoing evolutionary arms race that would manifests itself as diversifying selection across taxa, specifically on genes involved in host defense (whereas adaptations shared across the genus would be very difficult to detect). To test this, we reanalyzed microarray data comparing the transcriptional response of vervets and macaques to infection with SIV^33,34^. Unlike vervets, macaques are not natural hosts of SIV and generally develop AIDS-like symptoms upon infection. If some of the selection signals reflect adaptation to SIV in vervets, we would expect selection scores to be elevated in genes that are differentially expressed in vervets — but not in macaque — as a response to infection. Indeed, selection scores are much higher in genes that show a significant expression difference before and after infection in vervets only, as compared to genes showing an expression difference in both species (one-sided Mann-Whitney U test p<10^−7^) or in macaque only (p<10^−4^) (Supplementary Fig. 26). Conversely, vervet-specific (but not shared or macaque-specific) differentially expressed genes are significantly enriched in high selection scores (p<0.003, p>0.99 and p>0.99, respectively). To further investigate the underlying mechanisms, we grouped differentially expressed genes by coexpression pattern using weighted gene co-expression network analysis (WGCNA, Supplementary Figs. 27 and 28)^35^. Five out of the 33 gene co-expression modules show a significant enrichment for genes with high selection scores (FWER<0.05; Fig. 3C, Supplementary Fig. 28, Supplementary Data 6). Remarkably, the significant modules share similar expression patterns with strong changes in vervets post-infection and very weak, mostly opposing, signals in macaques. In particular, all the modules that are enriched for diversifying selection show changes in gene expression in vervet six days postinfection, which is around the time that the virus becomes detectable and activates early immune responses. Two modules also show expression differences in the chronic stage (115 days post infection), which is most relevant for progression to immunodeficiency. We ran GO-enrichment analysis separately on the genes in the enriched WCGNA modules showing early (Fig. 3C, top three panels) and late expression changes (Fig. 3C, bottom two panels). We found 32 and 20 significantly enriched GO-categories, respectively, many of which are involved in response to HIV in humans (Supplementary Data 7 and 8). For example, for early expression response, enriched GO-categories include *clathrin-mediated endocytosis^36^*, *autophagosome assembly*, *positive regulation of type I interferon (IFN-I) production^33,34^* and *innate immune response*. This is consistent with recent findings that in macaques the IFN-I response is delayed in response to SIV infection and inhibited during the first week of SIV infection^37^, while natural hosts mount a very early and transient IFN-I response^33,34^. Conversely, the three most highly enriched GO-categories for genes in modules with late expression changes are *positive regulation of natural killer (NK) cell activation*, *regulation of cellular response to heat^38^*, and *somatic hypermutation of immunoglobulin genes^39^*, consistent with differences in NK cell responses during SIV infection in natural hosts as compared to non-natural hosts, the lack of viral replication in B cell follicles (Tfh cells) and preservation of lymph nodes immune function in natural hosts in contrast to macaques, as well as a better adaptation to the stress induced by the chronic infection^40–43^.

While enrichment analysis identifies categories of genes under selection, and is likely driven by large numbers of genes with moderate effects, the highest selection scores identify candidate regions for strong selection (Fig. 4, Supplementary Data 9). The highest score is for an uncharacterized gene on chromosome 6 (Fig. 4B) with 97% sequence identity to human RAN binding protein 3 (RANBP3), a gene connected to influenza A virus replication^44^ and that is involved in nucleocytoplasmatic export of RNAs from human T-cell leukemia virus type 1 (HTLV-I)^45^ and HIV^46,47^. The gene with the second highest score is NFIX nuclear factor I/X (Fig. 4D), a transcription factor that binds the palindromic sequence 5’-TTGGCNNNNNGCCAA-3 in viral and cellular promoters. Nuclear factor I proteins can serve as a transcription selectivity factor for RNA polymerase II, and play a critical role in transcription and regulation of JC virus in humans^48^ and Simian virus 40 in vervet cells^49^. Remarkably, these closely related viruses are usually harmless but cause disease under immunodeficiency, specifically in SIV/HIV infection in macaque^50^ and human^51^. However, the lack of common genetic variants in the coding sequence of this gene, suggests that selection is more likely to have targeted regulatory variants. See Fig. 4 and Supplementary Data 9 for further candidates.

**Fig. 4:**
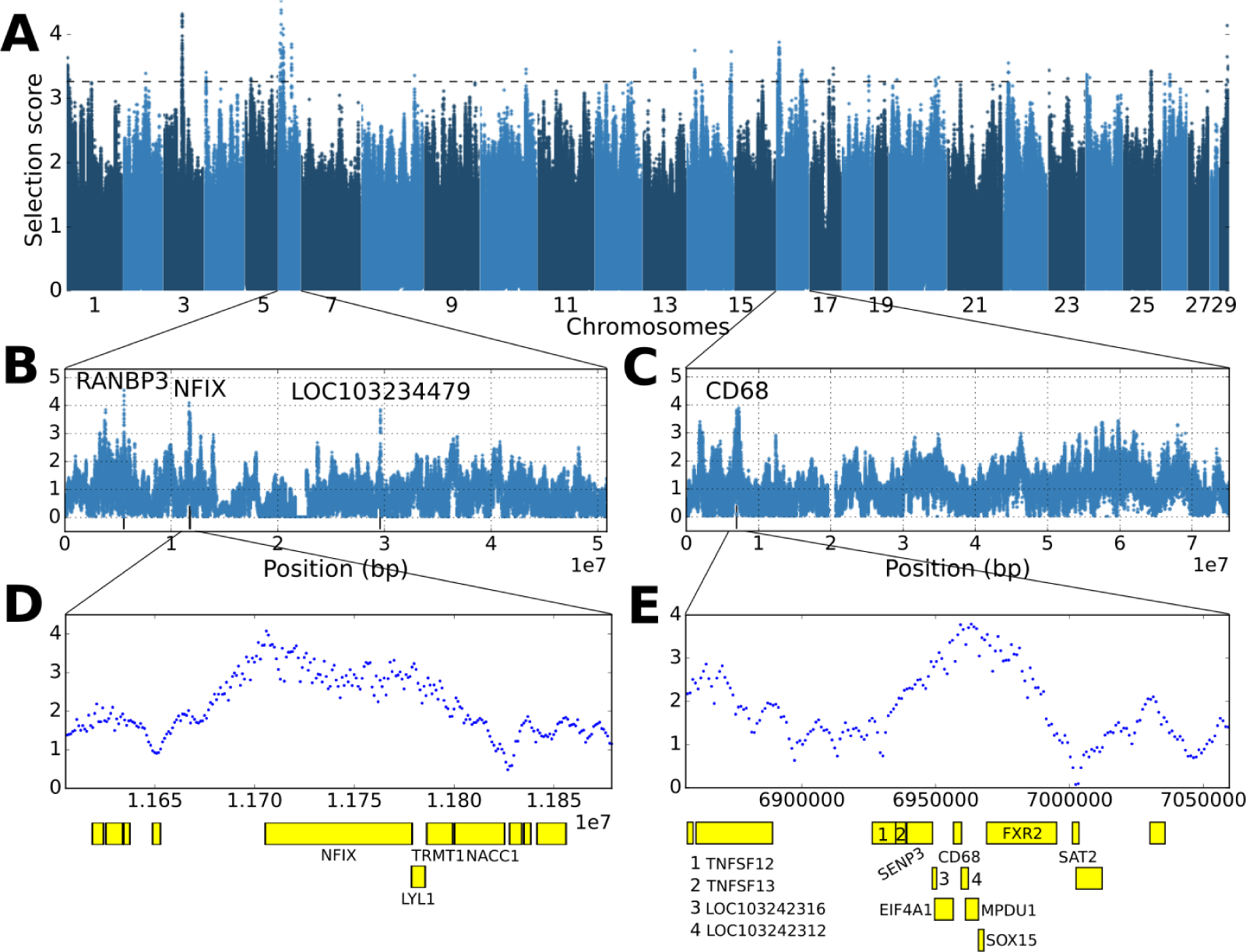
Selection scores across the genome and candidate genes with strong selection signals. (A) Manhattan plot of selection scores across all chromosomes. (B-C) Selection scores along chromosomes 6 and 16, respectively (D) Magnification of the region containing NFIX. (E) Magnification of a peak containing multiple candidates (Supplementary Data 9), among them CD68 (Cluster of Differentiation 68), a glycoprotein highly expressed on monocytes/macrophages, and FXR2 (Fragile X mental retardation, autosomal homolog 2) that interacts with HIV-1 Tat gene. Slightly downstream of the shown region, we note the highly scoring gene KDM6B (lysine-specific demethylase 6B), which is upregulated by HIV-1 gp120 in human B-cells.

In conclusion, we genetically characterized the entire vervet monkey genus, *Chlorocebus*. With some important exceptions, our analysis supports the existing classification into taxa, but suggests that their origin cannot readily be described using a simple phylogenetic tree. A screen for diversifying selection across taxa reveals a strong excess of signals in genes that interact with viruses, and we present evidence that part of the signal results from vervet co-evolution with SIV. Interestingly, the genes identified do not include the virus (co-)receptor genes involved in the virus docking mechanism, but rather genes involved in cell signalling and transcriptional regulation, consistent with recent results suggesting that natural selection has shaped primate CD4+ T-cell transcription^52^, and suggesting adaptation to living with the virus rather than avoidance of infection. Indeed, one of the highest scoring genes controls the expression of a virus known to cause disease under SIV-induced immunodeficiency. Along with the genomic polymorphism data, we provide a catalogue of candidate genes and genomic regions that could prove useful in the quest for antiviral vaccinations and therapies.

## Methods

### Sequencing, variant detection and filtering

Samples were sequenced at variable coverage (Illumina 100bp paired-end, Supplementary Data 1) with a total coverage of 798X and a median coverage of 4.4X. Sequences were aligned against the ChlSab1.1 reference^9^ using bwa-mem^53^. On average, more than 98% of the reads mapped for all taxa, suggesting that reference bias is weak. Following the GATK recommended workflow^54,55^, alignments were locally realigned, base quality scores were recalibrated using a first round of variant calling and variants were detected using GATK UnifiedGenotyper. Biallelic SNP calls were hard filtered with a combination of GATK best practises^55^ and custom filters to yield the data set used for further analysis (Supplementary Table 1, Supplementary Fig. 1). Note that our filtering was optimised to minimise bias rather than false positive rate. Given the large differences in coverage between individuals (2X-45X), a stringent control on false positive rate would lead to a bias towards lower diversity (and especially a lower number of singletons) in low coverage samples. We suggest that for population genomic analysis it is conservative to reduce bias at the cost of increased noise. The ancestral state for each SNP was determined by aligning the macaque reference genome, rheMac2, against ChlSab1.1, only using one-to-one mappings with a minimum length of 200bp.

### Accessible genome size

To compare levels of polymorphism and divergence across individuals and to previous studies, we measured the proportion of the genome accessible to our variant detection process. In particular, we excluded all sites that did not pass our quality filters, and, for each individual, all sites for which UnifiedGenotyper could not make a genotype call (Ns) (Supplementary Fig. 2).

### Diversity and divergence

Nucleotide diversity was calculated by computing the number of pairwise differences for each comparison divided by the accessible genome size for each pair as derived above. For each group, nucleotide diversity was estimated as the average of within-group comparisons. We found 16-27 million SNPs segregating within species, corresponding to an average number of pairwise differences per site (nucleotide diversity) of 0.17-0.22% (Fig. 1C, Supplementary Fig. 5) and effective population sizes generally above 35,000 (except for *aethiops* for which we estimate ~29,000). Site frequency spectra within taxa (Supplementary Fig. 3) generally agree with neutral expectations, except for a general lack of low frequency variants and an excess of high frequency derived variants, most likely as a consequence of low power to call low frequency variants and erroneous inference of the ancestral state, respectively.

Divergence was calculated as the average of pairwise differences across all comparisons of two groups. Two taxon site frequency spectra (Supplementary Fig. 4) generally show both, private variation, fixed differences and shared variation, except for *hilgerty/cynosorus/pygerythrus* which show few fixed differences and highly correlated allele frequencies.

To assess the relative contribution of geography and taxons label to explaining the genetic relatedness among vervets, we calculated principal components (PCs) from autosomal SNPs using PCAdapt version 05/26/14 in mode *fast^56^* setting K=6, which gave the best fit to our data. Next, for each of the first six PCs we performed likelihood ratio tests in R (function anova with option test=’LRT’) to test whether a linear model “PC ~ latitude + longitude + taxon label” gave a significantly better fit than a model using either only geography or only taxon (Supplementary Table 3).

F_ST_ was calculated using the Weir-Cockerham estimator^57^ using vcftool^58^ for all autosomal SNPs. For each pairwise comparison, we summarised FST-values in minor allele frequency (MAF) bins (Supplementary Fig. 6; the maximum across MAFs in Fig. 1C).

### SIVagm phylogenetic analyses

A 602 bp pol integrase fragment of SIVagm obtained as described previously^10^ was used for phylogenetic analyses of a large sample of SIVagm strains from the different subtaxa of vervet with different origin. pol nucleotide sequence alignments were obtained from the Los Alamos National Laboratory HIV Sequence Database (http://hiv-web.lanl.gov). Newly derived SIV sequences were aligned using MUSCLE^59^ and alignments were edited manually where necessary. Regions of ambiguous alignment and all gap-containing sites were excluded.

Phylogenetic trees were inferred from the nucleotide sequence alignments by the neighbor-joining method using the HKY85 model of nucleotide substitution^59,60^ implemented using PAUP*^61^. The reliability of branching order was assessed by performing 1,000 bootstrap replicates, again using neighbor-joining and the HKY85 model. Phylogenetic trees were also inferred by maximum-likelihood using PAUP* with models inferred from the alignment using Modeltest^62^. The neighbor-joining tree topology was used as the starting tree in a heuristic search using TBR branch swapping.

### Admixture analysis

The software Admixture 1.23^17^ was run on autosomal SNPs filtered for minor allele frequency > 5%, converted to binary .bed format using GATK VariantToBinaryPed and LD pruned using the plink flag *‐‐indep 50 10 2* (Fig. 3A and Supplementary Fig. 8). Cross-validation was run to get an indication of an appropriate number of ancestral genetic clusters, K (Supplementary Fig. 7). Cross-validation error was low between K=3 and K=7 with a global minimum at K=4 and a local minimum at K=6.

### Cross-taxon gene flow across time

Missing genotypes in our autosomal SNP calls were imputed using beagle 4^63^, version 03Oct15.284:

~~~
java ‐jar beagle.jar gl=biallelic_pass_snps.vcf out=beagle_out.vcf ibd=false
~~~

Samples were phased using shapeit.v2.r837.GLIBCv2.12.Linux^64^:

~~~
shapeit ‐phase ‐‐input-vcf beagle_out.vcf ‐‐window 0.1 ‐O phased.tmp
~~~

~~~
shapeit ‐output-vcf ‐‐input-haps phased.tmp ‐O phased.vcf
~~~

For each geographic sample group of interest, we chose the individual with the highest coverage for further analysis. For pairs of individuals from different groups, we extracted the alleles that segregate within or between the two individuals and their phase as needed as input to MSMC. The number of informative sites between two segregating variants was determined for each pair of individuals separately from the all-sites VCF (the whole genome including non-variant sites) by counting the number of non-filtered sites for which both individuals had genotype calls.

Three runs of MSMC2 (https://github.com/stschiff/msmc2) were produced for each pair, two inferring coalescent rates across time within each of the two samples

~~~
msmc2 ‐I 0,1 ‐o within_1 input_chrom1 … input_chrom29
~~~

~~~
msmc2 ‐I 2,3 ‐o within_2 input_chrom1 … input_chrom29
~~~

and one run for inferring coalescent rates across time between the two samples:

~~~
msmc2 ‐P 0,0,1,1 ‐o between input_chrom1 … input_chrom29
~~~

The outputs of these runs were combined by interpolating the mid-point of each time interval in the former two on the mid-points in the latter run. Cross-coalescent rate was calculated as 2^*^between/(within_1+within_2) (Figs. 2, B and C, Supplementary Figs. 9 and 10, Supplementary Note 1). Evolutionary time was scaled to years using a mutation rate of 1.5^^*^^10^−8^ and a generation time of 8.5 years.

### D-statistic

D-statistic was calculated from autosomal SNPs using Admixtools 3.0^19^, treating samples from each country as a populations and performing all tests that were consistent with the country UPGMA tree shown in Fig. 2D. Macaque was used as an outgroup and the analysis was restricted to sites where the macaque allele could be inferred. Due to limitations in Admixtools the analysis was restricted to vervet chromosomes 1-24. Block-jackknifing was performed with Admixtools standard settings.

### Diversifying selection scan

Autosomal biallelic PASS SNP genotypes were converted to XP-CLR input format. For each comparison of two groups, we excluded SNPs if they were not segregating within or between the groups or if they had more than 20% missing genotypes across the two groups. The genetic map from Huang et al.^65^ was interpolated to our SNP positions to get genetic distance in Morgan.

XP-CLR was run on all 30 possible comparison of the 6 African taxa (for each comparison, using each taxon once as objective and once as reference population). Parameters supplied to XP-CLR were *-w1 0.001 500 1000 ‐p0 0*, meaning that a set of grid points as the putative selected allele positions are placed along the chromosome with a spacing of 1 kb, the sliding window size was 0.1 cM around the grid points and if the number of SNPs within a window is beyond 500, some SNPs were randomly dropped to control for SNP number. Alleles were assumed unphased (*-p0*) and SNPs in high LD were not down-weighted (final 0).

To find loci that were repeatedly under diversifying selection across several group comparisons, we calculated for each grid point the root mean square selection score across all 30 comparisons (Fig. 4A). To test whether these scores capture biological signal, we confirmed that scores are significantly higher in genic (introns + exons) than in intergenic regions (Supplementary Fig. 20, one sided Mann-Whitney U test, p<10^−300^). Since the Mann-Whitney U test assumes independence of scores, which is not totally met due to linkage disequilibrium, we also calculated the average of the mean selection score across each gene and compared the resulting value to a background distribution. The background distribution was obtained by first concatenating all chromosomes in a circle and randomly shifting (rotating) the scores against their genomic positions, then calculating mean gene scores from these rotated data. We again found that genic scores are significantly larger (p<10^−5^).

### Gene set enrichment analysis

z-score transformed selection scores across genes (exons and introns) were used for gene enrichment analysis. Gene locations were extracted from NCBI *Chlorocebus sabaeus* Annotation Release 100 (http://www.ncbi.nlm.nih.gov/genome/annotation\_euk/Chlorocebus\_sabaeus/100/). To test for enrichment in gene ontology (GO) terms, we first used the the R-package TopGO^22^ with the “weight01” algorithm which allows to account for the hierarchical structure (and thus overlap) of GO terms when testing significance and thereby implicitly corrects for multiple testing. We restricted the analysis to 5777 GO terms with more than ten genes in our annotation. Note that our gene scores are not biased by gene length because we are calculating the average score across genes rather than taking the maximum score. However, enrichment results are qualitatively similar if the maximum is taken. Results are also similar if only exons are used (rather than exons + introns). We also note that the most significantly enriched categories contain many genes (Supplementary Data 4) and do not show strong clustering in particular genomic regions.

Negative selection on linked deleterious alleles (background selection) can produce genomic signatures similar to local adaptation. This is, because background selection locally decreases effective population size, which locally increases genetic drift within species and can thus contribute to increased allele frequency differentiation. To test whether our results could be dominated by signals of negative selection, we downloaded evolutionary conservation scores across 46 vertebrate species (phastCons46way, http://hgdownload.cse.ucsc.edu/goldenPath/hg19/phyloP46way/vertebrate/)^24^, and calculated mean conservation scores across genes, while excluding genes with less than 50% of their bases covered by scores. The resulting gene conservation scores (conscores) show weak but significant correlation with our selection scores (Pearson’s ?=0.11, p<10^−38^). We used these conscores to run the same GO enrichment analysis as for our selection scores using TopGO. Comparing the results, we note that GO enrichments are generally much less significant for conscores (46 vs. 167 scores significant at p<0.01, smallest p-value 1.1^*^10^−4^ vs. 4.2^*^10^−17^). There is some overlap in significantly enriched GO terms, most notably, *proximal/distal pattern formation, anterior/posterior pattern specification*, and *embryonic forelimb morphogenesis* are highly significant in both analysis (Supplementary Fig. S23), but most of the significant selection score enrichments are not significant using conscores. In particular, the highly significant and virus-related categories *positive/negative regulation of transcription of RNA polymerase II promoter*, and *viral process*, are not significant for conscores. This leads us to conclude that while our selection scan is likely to pick up signals of negative selection, in particular in developmental categories, there is no indication that virus-specific enrichments are driven by background selection.

### HIV-1 human interaction categories

The NCBI HIV-1 Human Interaction Database^27^ was downloaded from https://www.ncbi.nlm.nih.gov/refseq/HIVInteractions (last access Jannuary 2016). We only kept categories from the database that had ten or more genes in our annotation. We implemented the sumstat statistic^5^ to compare observed and expected gene-averaged selection scores in HIV-1 Human interaction categories to random sets of known genes. We found that 45 out of 166 categories show significant enrichment at Holm-Bonferroni FWER<0.05 (Fig 3B, Supplementary data 5).

### Gene expression analysis

We conducted Weighted Gene Co-expression Network Analysis (WGCNA) on gene expression data of vervets and macaques essayed at different time points pre‐ and post-infection with SIVagm and SIVmac, respectively^33^, using the WGCNA R package as previously described^35,66^.

We used as a starting point the list of genes with differentially expressed transcripts in CD4+ cells before and after SIV infection in either vervet or rhesus^33^. Correlation coefficients were constructed between expression levels of genes, and a connectivity measure (topological overlap, TO) was calculated for each gene by summing the connection strength with other genes. Genes were then clustered based on their TO, and groups of coexpressed genes (modules) were identified. Each module was assigned a color, and the first principal component (eigengene) of a module was extracted from the module and considered to be representative of the gene expression profiles in a module. We identified 36 modules (Supplementary Figs. 27 and 28, Supplementary Data 6).

## Acknowledgments

Samples were collected through the UCLA Systems Biology Sample Repository funded by NIH grants R01RR016300 and R01OD010980 to N.F. For permits allowing us to collect samples, we thank the Gambia Department of Parks & Wildlife Management; Botswana Ministry of Environment & Wildlife and Tourism; Ghana Wildlife Division, Forestry Commission; Zambia Wildlife Authority; Ethiopian Wildlife Conservation Authority; Ministry of Forestry & the Environment, Department of Environmental Affairs, South Africa; Department of Economic Development and Environmental Affairs, Eastern Cape; Department of Tourism, Environmental and Economic Affairs, Free State Province; the Ezemvelo KZN Wildlife in KwaZulu Natal Province; and the Department of Economic Development, Environment and Tourism, Limpopo Province. We also thank Dr. Gene Redmond and St. Kitts Biomedical Research Foundation, and Dr. Martin Antonio and Medical Research Council (MRC) The Gambia Unit for facilitating sample collection in St. Kitts and Nevis and the Gambia, respectively. We thank Drs. Jason Brenchley, Keith Reimann (R24OD010976) and Jean Baulu and the Barbados Primate Research Center and Wildlife Reserve for providing samples from Tanzania origin and Barbados vervets. For help with sample collection we thank Drs. Trudy Turner, Christopher Schmitt, Jennifer Danzy-Cramer and J. Paul Grobler, and Ms. Yoon Jung, and Mr. Oliver Pess Morton. H.S. has been supported by a travel grant of the Austrian Ministry of Science and Research. We thank Reena Halai for help with figure design. CA is supported by RO1 AI119346 from NIAID. We acknowledge the support of the NINDS Informatics Center for Neurogenetics and Neurogenomics (P30 NS062691). We would like to thank Ms. Fuying Gao for assistance with microarray data analysis. All genomic data for the sequenced vervet subspecies in this study are available through the NCBI SRA public repositories under NCBI BioProject numbers PRJNA168521, PRJNA168472, PRJNA168520, PRJNA168527, PRJNA168522. Variant call format (VCF) files will be uploaded for public access.

